# Alirocumab attenuated plaque inflammation and PCSK9-induced proinflammatory signalling in M1 macrophages independently of lipid lowering

**DOI:** 10.1101/2025.11.20.689631

**Authors:** C. Espadas, M. Soto-Catalán, M. Romero-Cote, M. Kavanagh, I. Herrero-Del Real, A. Ortega-Hernández, J. Lumpuy-Castillo, D. Gómez-Garre, J. Egido, J. Tuñón, C Gómez-Guerrero, Ó. Lorenzo

## Abstract

**Background:** Proprotein Convertase Subtilisin/Kexin Type 9 (PCSK9) may directly promote vascular inflammation beyond its action on LDL-C degradation. We investigated whether PCSK9 may exacerbate proinflammatory signalling of M1 macrophages and if alirocumab could attenuate this effect and plaque progression by LDL-C independent mechanisms.

**Methods:** ApoE⁻/⁻ mice were treated with alirocumab for 13 weeks and aortic arches were isolated for atherosclerotic plaques characterization based on lesion size and lipid and macrophage infiltration. Also, plasma and splenic monocytes/macrophages were assessed by flow cytometry, and PCSK9, the lipid profile, inflammatory cytokines were measured by qPCR or western blot. Cultured THP-1-derived M1 macrophages were stimulated with PCSK9 and evaluated for TLR4-NFκB-NLRP3 activation and cytokine production. PCSK9 and proinflammatory factors were analysed in 1190 patients with acute coronary syndrome (ACS).

**Results:** Alirocumab reduced plaque lesion (0.42-fold; p< 0.05) and lipid (0.63-fold; p< 0.01) and macrophage (0.61-fold; p<0.05) infiltration, mainly the M1 subtype (0.37-fold; p<0.01), as well as TLR4, NLRP3 and caspase-1 expression (0.49-fold, 0.51-fold and 0.51-fold, respectively; p<0.05), without altering LDL-C. Also, it decreased proinflammatory cytokines but enhanced anti-inflammatory factors and M2 markers at the descending aorta. Alirocumab enriched circulating Ly6C^low^ monocytes (1.51-fold; p<0.05) and splenic M2 macrophages (1.32-fold; p<0.01), while reducing M1 (0.62-fold; p<0.05). In cultured M1 macrophages, PCSK9 overexpressed proinflammatory cytokines (i.e., CXCL9, CXCL10, TNF-α, IL-1β, IL-6) and downregulated anti-inflammatory mediators (i.e., CCL17, TGM2, TGF-β1, IL-10), promoted NFκB-p65 nuclear translocation, and NLRP3 and gasdermin-D activation. However, TLR4 inhibition or silencing blunted these effects. In addition, there was a positive association between PCSK9 with hsCRP and FGF-23 plasma levels in ACS patients.

**Conclusions:** PCSK9 may be released in parallel to proinflammatory factors such as hsCRP and FGF-23 in ACS patients. However, alirocumab could exert lipid-independent anti-inflammatory effects through activation of the TLR4-NFκB-NLRP3 signaling in M1 macrophages, promotion of M2, and reducing plaque inflammation.

## 1. Introduction

Atherosclerosis is a chronic inflammatory disease initiated and sustained by the accumulation of lipids, particularly oxidized low-density lipoproteins within the artery wall, which triggers complex inflammatory responses involving vascular and immune cells^1^. Particularly, monocytes drive atherogenesis by infiltrating into atherosclerotic plaques, differentiating into macrophages and, upon lipid uptake, becoming foam cells that amplify inflammation and plaque growth. Moreover, macrophages can polarize to proinflammatory M1 or tissue-repair M2 macrophages in response to multiple signals from microenvironment^2^. Activation of NFκB, primarily through Toll-like receptors (TLR), can be a crucial factor promoting the M1 phenotype (suppressing M2 polarization) to overexpress proinflammatory cytokines, intensifying endothelial activation and monocyte recruitment^3^. Also, activation of the NLRP3 pathway can lead to upregulation of M1-associated cytokines, such as IL-1β and CXCL10, and decrease of M2-associated factors like ArgI and CD206^4^. Thus, despite lipid accumulation remains a key driver of plaque formation, the persistent inflammatory milieu may further increase residual cardiovascular risk^5^. In this sense, new anti-atherosclerotic therapies could address vascular inflammation to protect from cardiovascular injuries.

Traditional anti-atherosclerosis drugs, particularly statins, possess hypolipidemic and anti-inflammatory properties^6^. However, they can also interact with other medications and evidence regarding their direct impact on the inflammatory system, independent of lipid reduction, remains unclear. In this line, the pro-protein convertase subtilisin/kexin type 9 (PCSK9) is highly expressed in hepatocytes and has been shown to induce low-density lipoprotein (LDL)-receptor degradation, reducing the LDLR-mediated clearance of pathogenic lipids^7^. Interestingly, PCSK9 could be also upregulated in infiltrated macrophages leading to local inflammatory responses^8^. In fact, PCSK9 inhibitors such as alirocumab reduce plasma LDL-C and plaque progression^9^, but their potential direct anti-inflammatory actions has not been completely unveiled. Thus, we aim to investigate whether PCSK9 might play a role in the proinflammatory response by exacerbating the proinflammatory signalling of M1 macrophages through the TLR/NFκB/NLRP3 axe, and if alirocumab could confer anti-inflammatory effects independent of LDL-C reduction.

## 2. Methods

### 2.1 Human study

We analysed 1,190 patients with acute coronary syndrome (ACS) with/without ST elevation of the BACS & BAMI (Biomarkers in ACS & Biomarkers in Acute Myocardial Infarction) study carried out in five hospitals in Madrid. Inclusion and exclusion criteria were previously described^10^. On admission, clinical variables were recorded, and 12h-fasting blood was taken for analysis. Then, EDTA-plasma samples were immediately isolated by blood centrifugation at 2500 rpm (20 min) at 4°C and stored at −80°C until use. PCSK9 was determined using enzyme-linked immunosorbent assay (ELISA) with anti-PCSK9-specific antibody (ELLA kit SPCKB-PS00321, ProteinSimple, Bio-Techne). Also, a specific panel of proinflammatory factors composed by high-sensitivity C-reactive protein (hs-CRP), monocyte chemoattractant protein-1 (MCP-1), fibroblast growth factor-23 (FGF-23), and interleukin-18 (IL-18) was quantified using the latex-enhanced immunoturbidimetry (ADVIA 2400 Chemistry System), the ELISA kits (DCP00 human MCP-1 from R&D Systems, and human FGF23-C-Term from Immutopics Inc), and the ELLA kit SPCKB-PS000501 for IL-18.

The research protocol conformed to the ethical guidelines of the 1975 Declaration of Helsinki as reflected by approval of the human research committees of the institutions participating in this study (Ref.: 05-07; 24 April 2007). An informed written consent was given to be signed by patients prior to the inclusion in the study.

### 2.2 *In vivo* studies

#### Mouse model of atherosclerosis

Apolipoprotein E-deficient (ApoE⁻/⁻) mice (B6.129P2-Apoe^tm1Unc^/J; The Jackson Laboratory, stock no. 002052, RRID:IMSR_JAX:002052), congenic on C57BL/6J after >10 backcross generations, were used as a model of hypercholesterolemia and accelerated atherosclerosis due to impaired ApoB- and ApoE-rich lipoprotein clearance, resulting in inefficient cholesterol removal and elevated plasma LDL-cholesterol (LDL-C) levels^11^. We fed ApoE⁻/⁻ mice (males, 10-12 weeks old; n=16) a Western diet (42% fat, 0.21% cholesterol; ref: D12079, Ssniff Spezialdiäten, Germany) for 15 weeks to accelerate plaque progression and complexity, providing an experimental model that better resembles advanced human disease^12^. Animals were housed in groups of 4-5 per ventilated cage (20–22 °C) and maintained on a 12h light/dark cycle, with ad libitum access to food and water. At the 2^nd^ week of Western diet, animals were randomly assigned to a control group (n=9) or to treatment group (n=7) with PCSK9 monoclonal antibody, alirocumab (10 mg/kg/week divided in two s.c. injections) as previously described^13^. Then, at the 15^th^ week, animals were euthanized under anaesthesia (ketamine, 100 mg/kg, and xylazine, 10 mg/kg) administered via intraperitoneal injection and total peripheral blood, aortic arch, the descending aorta, spleen, and liver were collected. ApoE⁻/⁻ mice represent an ideal model to investigate potential anti-inflammatory effects of PCSK9 inhibitors in atherosclerosis, independently of the lipid-lowering actions, since absence of ApoE precludes the hepatic clearance of remnant lipoproteins including LDL-C^14,15^. All procedures were approved by the FISS-FJD Animal Experimentation Ethics Committee and the Regional Government (Ref. PROEX 128.4-23) in accordance with EU Directive 2010/63/EU and national regulations (RD 53/2013).

#### Anthropometric, biochemical, and cytokine measurements

The animal body weight was weekly recorded. None of mice show infectious events along the model, and no differences were observed in the food and water intake. Plasma was obtained by centrifuge-separation (20 min, 2500 rpm, 4°C) from peripheral blood in EDTA containing tubes. The lipid profile was assessed using specific commercial kits for total cholesterol (TC) (Cell Biolabs, #STA-390), triglycerides (TG) (Cayman Chemical, #10010303), and LDL-C (CrystalChem, #79980). Plasma PCSK9 was evaluated by ELISA assay (R&D Systems, #MPC-900) whereas TNF-α and IL-6 were quantified using the Simple Plex™ ELLA automated immunoassay system (ProteinSimple, Bio-Techne). Moreover, liver PCSK9 expression was studied by western blot using and anti-PCSK9 (D7U6L) rabbit monoclonal primary antibody (Cell Signaling, #85813), as described below.

#### Quantification of atherosclerotic lesions and macrophage infiltration

During sacrifice, mice were perfused from heart with 10 ml cold PBS and entire aorta was collected. The aortic arch was embedded in OCT compound (Sigma-Aldrich, #6502) and stored at –80 °C. The aortic area of atherosclerotic plaque (μm2) was quantified from 1000 μm of the cardiac valve leaflets in serial sections of 7 μm obtained by cryostat (Leica CM1520). Sections were fixed with cold acetone (10 min, −20°C), washed (60% isopropanol) and stained for one hour with 2.5 mg/ml oil-red-O (ORO) (Sigma Aldrich, #O1391) and haematoxylin. The major lesion of atherosclerotic plaque was established in those sections with the maximal stained area. This region was quantified as the ratio of lesion area by the total aortic surface under microscopy (Zeiss Axioscope 5, Zeiss). In addition, the neutral lipid content was also quantified by ORO/haematoxylin within these sections.

Total macrophage infiltration at the atheroma plaques was evaluated by incubating OCT cryosections (just adjacent to maximal atherosclerotic lesion) with an anti-CD68 antibody (Abcam, #125212). First, sections were fixed in cold acetone, and endogenous peroxidase was quenched with methanol. Non-specific protein binding was blocked with goat serum (1h, room temperature) and primary anti-CD68 antibody (1:400) was incubated overnight at 4°C, followed by a biotinylated anti-rabbit secondary antibody (1:500, 1h, room temperature, Fisher Scientific, #11859200). The macrophage staining was developed by using the avidin–biotin complex (Vector Laboratories, #PK-7100), revealed with 3,3′-diaminobenzidine (DAB) (Abcam, #ab64238), and counterstained with haematoxylin. In all cases, for each mouse, two sections and four fields per section were analyzed with ImageJ software.

#### Characterization of macrophage infiltration in the aortic arch

Infiltrated macrophages were then characterized for M1 phenotype. Cryosections were fixed with 4% paraformaldehyde (PFA) for 10 minutes at room temperature, followed by blockade of non-specific binding using goat serum. Samples were then incubated overnight with anti-CD80 primary antibody (1:50, 4°C, Abcam, #ab254579) followed by a goat anti-rabbit IgG Alexa Fluor 488-conjugated secondary antibody (1:300, 1h, room temperature, ThemoFisher, #A-11008). Nuclear counterstaining was performed using DAPI for 10 minutes. After washing, sections were mounted and images were acquired using a Leica TCS SP5 confocal microscope (10× objective, 2.5× digital zoom). Fluorescence quantification was performed using ImageJ software.

Also, expression of TLR4, caspase-1 and NLRP3 were also analysed in these atherosclerotic lesions enriched in M1 macrophages (CD68^+^/CD80^+^) by immunohistochemistry with anti-TLR4 (Abcam, #218987), -caspase-1 (Santa Cruz, #sc-514) and -NLRP3 (Affinity-Bionova, #DF15549), respectively, at 1:100 dilution, followed by a biotinylated anti-rabbit secondary antibody (1:200, 1h, room temperature). Protein staining was revealed with the avidin–biotin complex and DAB, as above.

#### Aortic gene expression

Descending aorta was disaggregated using 1 mm diameter zirconium beads and a homogenizer (Bullet Blender Homogenizer, Next Advance Inc., Troy, NY, USA). Total RNA was isolated using TRIzol reagent (ThermoFisher Scientific, #15596026) according to the manufacturer’s protocol. RNA concentration and purity were assessed spectrophotometrically (Nanophotometer® N60, IMPLEN). The cDNA synthesis was performed using the High-Capacity cDNA Reverse Transcription Kit (ThermoFisher Scientific, #4368813) on a Veriti Thermal Cycler. Quantitative real-time PCR (qPCR) was carried out in 10 µL reactions containing cDNA, TaqMan™ Universal PCR Master Mix (ThermoFisher Scientific, #4318157), and gene-specific TaqMan assays *CXCL10* (Mm00445235_m1), *TNF-α* (Mm00443258_m1), *IL-1β* (Mm00434228_m1), *MCP-1* (Mm00441242_m1), *IL-6* (Mm00446190_m1), *TGF-β1* (Mm01178820_m1), *IL-10* (Mm00439614_m1), *CD163* (Mm00474091_m1), *ArgII* (Mm00477592_m1), and *CD206* (Mm01329359_m1)] on a StepOnePlus™ Real-Time PCR System (ThermoFisher Scientific). All samples were run in triplicate and those with cycle threshold (C_T_) variation exceeding 0.3 were excluded. The relative gene expression was calculated using the comparative ΔΔC_T_ method with *β-actin* (Mm02619580_g1) serving as the endogenous control.

#### Flow cytometry analysis

Characterization of monocyte and macrophage populations was achieved in peripheral blood and spleen. 200 µL of whole EDTA-blood was aliquoted and stabilized with Transfix Bulk (CytoMark, 300K Solutions, #TFB-20-1), according to the manufacturer’s instructions. Specific antibodies against monocyte markers (see below) were added to the samples and incubated for 30 min at 4°C. Then, the BD lysis buffer (BD Bioscience, #349202) was added for 10 min and samples were vortexed to lyse red blood cells, which were removed after centrifugation. The pellet with stained cells was washed and resuspended in cold PBS. On the other hand, spleens were harvested and kept in cold PBS to be mechanically disaggregated and filtered through 70 µm and 40 µm microfilters (Fisher Scientific, #10788201). Similarly, cell lysates were incubated with specific antibodies and fixed with 1% PFA in PBS.

For both blood and spleen samples, we tested a comprehensive panel of fluorochrome-conjugated antibodies targeting key monocyte and macrophage markers (BioLegend). We used: APC/Cyanine7 anti-mouse CD45 (clone 30-F11, #103116), APC anti-mouse/human CD11b (clone M1/70, #101212), FITC anti-mouse lymphocyte antigen 6 complex at locus C (Ly6C, clone HK1.4, #128006), PE/Cyanine7 anti-mouse F4-80 recombinant antibody (clone QA17A29, #157308), PE anti-Nos2 (iNOS) (clone W16030C, #696806), and Brilliant Violet 421™ anti-mouse CD206 (MMR) (clone C068C2, #141717). Moreover, CD45⁺/CD11b⁺-monocytes were further classified by the presence of Ly6C. Those Ly6C^high^ monocytes express higher levels of Ly6C and are mostly involved in proinflammatory activities, whereas Ly6C^low^ (lower Ly6C) correspond to monocytes involved in tissue repair and anti-inflammation^16^. Similarly, CD45⁺/F4-80⁺-macrophages were classified according to their polarization state as proinflammatory (M1) or anti-inflammatory (M2) macrophages. M1 macrophages were identified by the expression of inducible nitric oxide synthase (iNOS), while M2 macrophages were defined by the expression of the macrophage mannose receptor, CD206^17^. Then, flow cytometric acquisition was performed on a Cytoflex (Beckman Coulter) acquiring 10000 live events from peripheral blood and 50000 live events from spleen suspensions. Data analysis was conducted using CytExpert 2.3 and Kaluza Analysis Software (Beckman Coulter).

### 2.3 *In vitro* studies

#### Monocytes culture, differentiation, and polarization

The human THP-1 cell line of monocytes were donated by Dr. Martín-Ventura and cultured in Roswell Park Memorial Institute medium (RPMI 1640) supplemented with glutamine 2,05 mM (Gibco, #61870036), 10% heat-inactivated foetal bovine serum (Sigma-Aldrich, #F7524), and 1% Penicillin-Streptomycin (Sigma-Aldrich, #P0781), and maintained at 37°C and 5% CO_2_ in a humidified incubator. Monocytes were counted, seeded (1,5 × 10^6^ cells/p60-plate) and differentiated into naïve macrophages (M0) by adding 100 nM PMA (phorbol 12-myristate 13-acetate, Sigma, #P8139) for 72h. Differentiated macrophages were washed and primed for 48h with M1-polarization medium composed by 20 ng/ml IFN-γ (PeProtech, #300-02) and 10 pg/ml LPS (Sigma-Aldrich, #L3012) dissolved in RPMI-1640 (0,5% FBS) to polarize to the M1 phenotype^18,19^. M1 macrophages were confirmed by assessing the increased cytokine CXCL9/CCL17 ratio, as previously shown^20,21^.

#### Stimulation of M1 macrophages

M1 macrophages were stimulated for 3-24h with 2,5 ug/mL human recombinant PCSK9 (hPCSK9, Sigma-Aldrich, #SRP6285) in M1-polarization medium, as previously described^8^. TNF-α (100 ng/mL; PeProtech, #300-01A) or LPS (100 ug/mL; Sigma-Aldrich, #L3012) were used as positive controls of pro-inflammation. Some cells were also pre-treated with parthenolide (1h, 10 µM; Sigma-Aldrich) as NFκB inhibitor preventing IκBα degradation, or with MCC950 (10 µM; InvivoGen) as selective NLRP3 inhibitor that block its oligomerization and caspase-1 activation. TAK-242 (10 µM; MedChemExpress) and alirocumab (1μM) were used as specific TLR4 and PCSK9 inhibitors, respectively. All compounds except alirocumab were dissolved in DMSO, and vehicle-treated cells served as controls.

#### TLR4-gene Silencing

Small interfering (si)RNA sequence for the human TLR4 gene was purchased from ThermoFisher Scientific (Assay ID: S14195, #4390824). M1 macrophages were transfected with siRNA-TLR4 dissolved in using Lipofectamine® RNAiMAX reagent following manufacturer’s instructions (Invitrogen, ThermoFisher #13778100). After 24h, cells were washed with PBS and stimulated with PCSK9 or LPS. Confirmation of gene and protein silencing was done by qPCR and western blot (Suppl. Fig. 1A).

#### Macrophages’ gene and protein expression

Total RNA was isolated from cultured M1 macrophages and converted to cDNA for quantification of relative gene expression by qPCR, as described above. The human taqman assay IDs were as follows: *CXCL9* (Hs00171065_m1), *CXCL10* (Hs00171042_m1), *IL-1β* (Hs01555410_m1), *TNF-α* (Hs00174128_m1), *IL-6* (Hs0174131_m1), *CCL17* (Hs00171074_m1), *TGM2* (Hs01096681_m1), *IL-10* (Hs00961622_m1), *TGF-β1* (Hs00998133_m1), and *TLR4* (Hs00152939_m1) serving *18S* rRNA (Hs99999901_s1) as the endogenous control.

Total proteins were also extracted from M1 macrophages by adding lysis buffer (Tris 50 mM, NaCl 0.15 M, EDTA 2.5 mM, TRITON X-100 0.2%, IGEPAL 0.3%). Protein concentration was determined with the Pierce™ BCA Protein Assay Kit (ThermoFisher Scientific, #23225). Then, 20 ug per sample were loaded on SDS-PAGE gels, separated by electrophoresis, and transferred onto PVDF membranes (ThermoFisher Scientific, #88520). Nonspecific binding was blocked with 5% skimmed milk in Tris Buffer Saline-Tween 20, and primary antibodies were used to quantify the protein levels: anti-Phospho-IκBα (Ser32) rabbit monoclonal (Cell Signaling), -IκB-αlpha (L35A5) mouse monoclonal (Cell Signaling), -Phospho-NFκB p65 (Ser536) rabbit monoclonal (Cell Signaling), -RELA/NFκB p65 (F-6) (Santa Cruz), -NLRP3 rabbit monoclonal (Cell Signaling), -Cleaved gasdermin-D rabbit monoclonal (Cell Signaling), and -TLR4 (Abcam, #218987), serving -GAPDH (Invitrogen) or -β-actin (Sigma-Aldrich) antibodies for endogenous control. Secondary antibodies were goat anti-Mouse IgG (H + L) and goat anti-Rabbit IgG (H + L) (ThermoFisher Scientific, #G21040 and #G21234, respectively). The antibody–antigen binding was detected by chemiluminescence (iBright750, Invitrogen) and quantified by the Quantity One 4.6.6 software (BioRad). All experiments were done at least three times.

#### Cell immunofluorescence

Nuclear translocation of NFκB-p65 was detected by immunofluorescence. THP-1 cells (6.0 × 10^4^ cells/well) were seeded in 8-well chamber slides, differentiated and polarized to M1, as described above. Cells were stimulated with PCSK9 for 30 minutes, washed and fixed with 4% PFA in PBS (10 min, 4°C) followed by blocking (1% BSA) and permeabilization (0.1% Triton X-100, 1h, room temperature). Then, cells were incubated with primary anti-p65 (SantaCruz, #sc-8008) (1:80) at 4°C overnight, and secondary anti-mouse AlexaFluor-488 conjugated secondary antibody (1:200 dilution, 1h, ThermoFisher, #A-11001). The cell nuclei were stained with DAPI, and images were obtained by confocal microscope (Leica TCS SP5). Quantification was performed using ImageJ, counting 30 cells per field within 5 different fields. All experiments were done at least three times.

### 2.4 Statistical analysis

Data distribution in the human study was assessed using the Kolmogorov–Smirnov test to evaluate normality. Variables with non-normal distribution were expressed as median (interquartile range) and compared using the Kruskal–Wallis test, followed by Dunn’s post hoc correction for multiple comparisons. A quantile regression model was also applied to explore the relationship between plasma PCSK9 levels and selected clinical variables across different quantiles (0.25, 0.5, 0.75) of the outcome distribution. Two-tailed p-value < 0.05 were considered statistically significant. Data from animals and cells are expressed as mean ± standard deviation (SD) unless otherwise indicated. Normality was assessed using the Shapiro–Wilk test. Comparisons between two groups were performed using unpaired two-tailed Student’s *t*-test. For comparisons involving more than two groups, one-way analysis of variance (ANOVA) followed by Dunnet or Tukey’s post-hoc test was used. Non-parametric data were analyzed using Mann–Whitney *U* or Kruskal–Wallis tests, as appropriate. A p value < 0.05 was considered statistically significant. Statistical analyses were performed using GraphPad Prism version 8.0.1 (GraphPad Software, San Diego, CA, USA).

## 3. Results

### 3.1 Overexpression of proinflammatory factors and PCSK9 in acute coronary syndrome

We analyzed the potential association of PCSK9 and proinflammatory cytokines including hsCRP, MCP-1, FGF-23 and IL-18 in plasma from 1190 patients with ACS^22^. Briefly, subjects were 61.6 years-old, 77.1% of them were men, showed a body mass index of 28.3 kg/m^2^, and exhibited hypertension (57.1%), dyslipemia (60.1%), and previous coronary artery disease (20.2%). Interestingly, by quantile regression we observed a significant association of PCSK9 with hsCRP (0.75 quantil) and FGF-23 (0.75 quantil), even after adjustment by sex, age, BMI, and additionally, dyslipidemia, previous statin use, and previous coronary artery disease (Figure 1A). Similarly, PCSK9 levels differed significantly across tertiles of both hsCRP and FGF-23. In fact, PCSK9 concentrations were higher in the third tertile compared to the first and second tertiles of both hsCRP and FGF-23 (Figure 1B). These findings demonstrate that higher hsCRP and FGF-23 levels can be accompanied by increased circulating concentrations of PCSK9 in patients with ACS, suggesting a lipid-independent pro-inflammatory role of PCSK9 in this disease.

**Figure 1.**
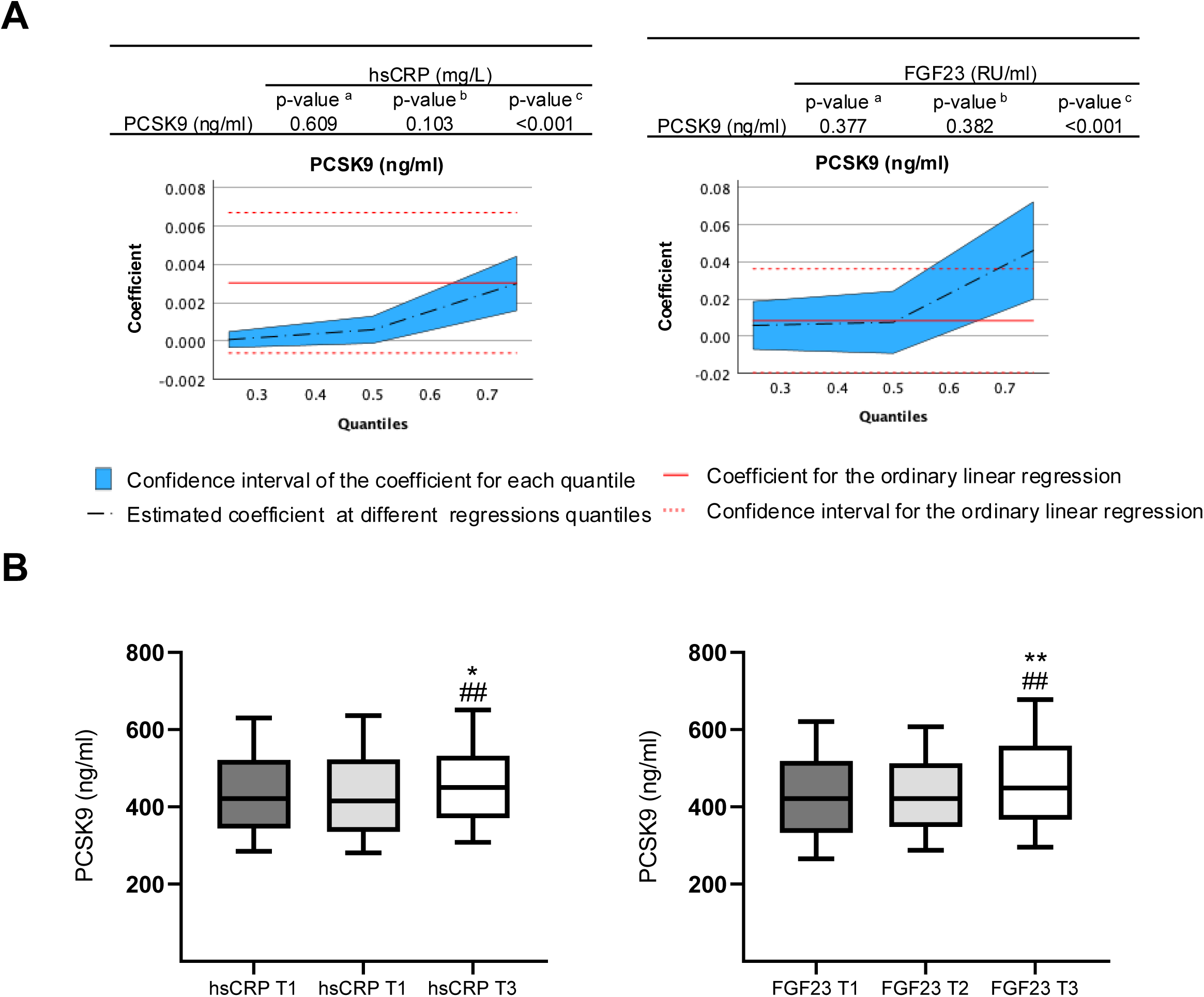
Plasma PCSK9 and proinflammatory factors after acute coronary syndrome. **(A)** Associations between PCSK9 with hsCRP and FGF23 plasma levels. Coefficients were estimated at the 0.25 (a), 0.50 (b), and 0.75 (c) quantiles of each factor by using quantile regression models adjusted for sex, age, BMI, dyslipidemia, statin use, and coronary artery disease. A p < 0.05 was considered significant. **(B)** Distribution of plasma PCSK9 levels by hsCRP and FGF-23 tertiles. Data are shown as box plots including median and 25^th^-75^th^ percentiles (whiskers represents the 10^th^-90^th^ percentiles). *p<0.05 and **p<0.01 vs. T1; ^#^p<0.05 and ^##^p<0.01 vs. T2.

### 3.2 Alirocumab reduced inflammation but not hyperlipidemia in ApoE⁻/⁻ mice

As previously described^12,23^, after 15 weeks of Western diet, ApoE⁻/⁻ mice showed higher levels of plasma lipids (total cholesterol, LDL-cholesterol and triglycerides) and increased levels of proinflammatory cytokines, TNF-α and IL-6 (Table 1). Interestingly, administration of the PCSK9 neutralizing antibody, alirocumab (10 mg/kg/week) for 13 weeks overexpressed liver and plasma PCSK9 (Suppl. Fig. 1B) as observed in human^24^, but did not modify the body weight and lipid profile (Table 1). However, alirocumab significantly attenuated serum levels of TNF-α and IL-6 (Table 1) and reduced the proinflammatory phenotype of circulating mononuclear cells. By flow cytometry, we observed an augmented number of anti-inflammatory CD45^+^/CD11^+^-Ly6C^low^ monocytes in alirocumab treated mice compared to untreated animals (5.6 ± 1.5% vs. 3.6 ± 1.3%, respectively; p<0.05), while the proinflammatory CD45^+^/CD11^+^-Ly6C^high^ monocytes were not altered (Figure 2A). In spleen, alirocumab also reduced the number of proinflammatory M1 (CD45^+^/F4-80^+^/iNOS^+^) macrophages (0.5 ± 0.2% vs. 0.8 ± 0.3%, respectively; p<0.05) and enhanced the M2 (CD45^+^/F4-80^+^/CD206^+^) macrophages (48.0 ± 4.5% vs. 36.3 ± 8.5%, respectively; p<0.01) (Figure 2B).

**Figure 2.**
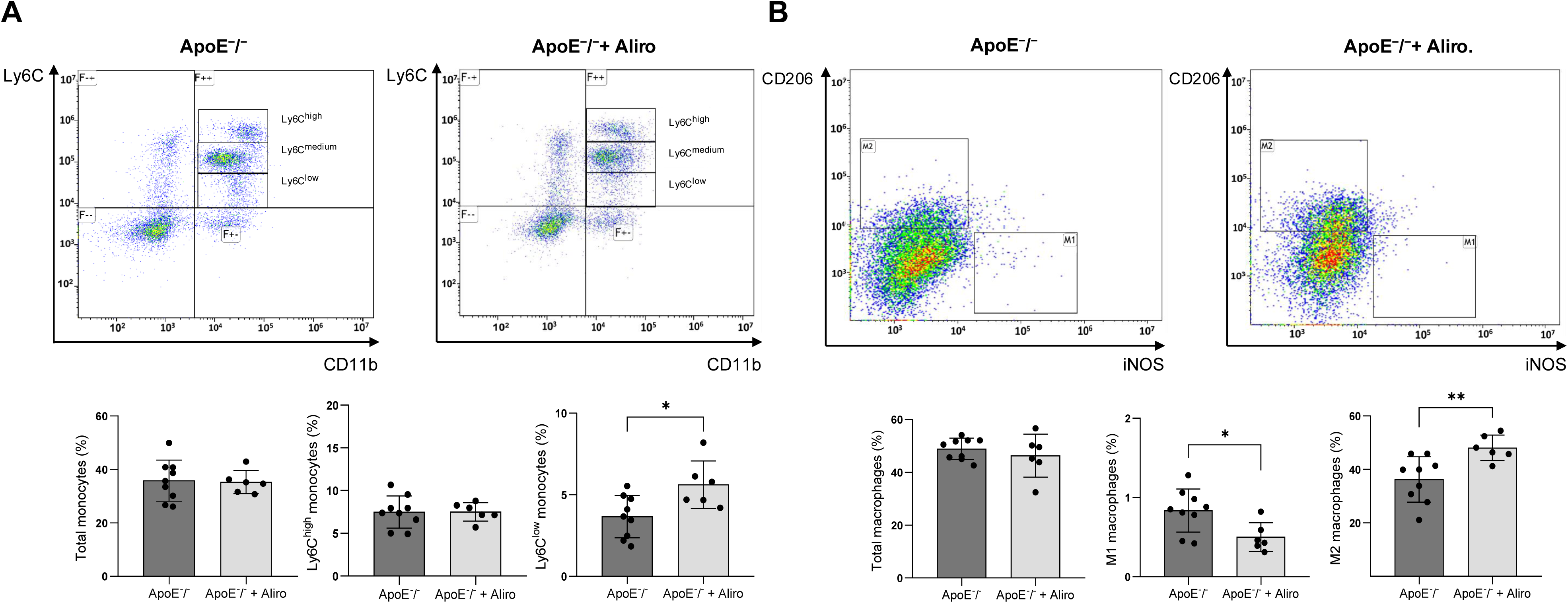
Alirocumab enhanced Ly6C^low^ monocytes and M2 macrophages in ApoE⁻/⁻ mice. **(A)** Top, gating-analysis for CD11b and Ly6C (high, medium, and low phenotypes) positive staining in plasma monocytes from ApoE⁻/⁻ and ApoE⁻/⁻ + alirocumab mice. Bottom, quantification of total, Ly6C^high^ and Ly6C^low^ monocytes. **(B)** Top, gating-analysis for iNOS and CD206 positive staining in spleen macrophages from ApoE⁻/⁻ and ApoE⁻/⁻ + alirocumab mice. Bottom, quantification of total, M1 and M2 macrophages. *p<0.05 and **p<0.01 vs. ApoE⁻/⁻ mice, by unpaired two-tailed Student’s *t*-test. N=6-9, per group.

**Table 1.**
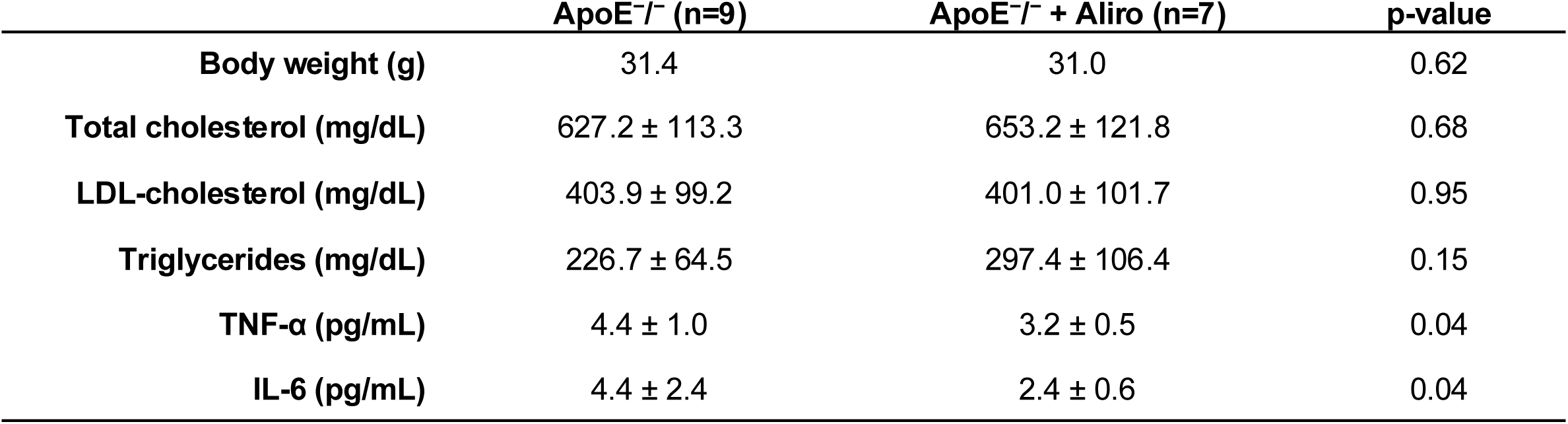
Alirocumab reduced plasma cytokines in ApoE⁻/⁻-Western diet-treated mice. Body weight, plasma lipid profile (total cholesterol, LDL-C and triglycerides), and circulating proinflammatory cytokines (TNF-α and IL-6).

|  | **ApoE<sup>-/-</sup> (n=9)** | **ApoE<sup>-/-</sup> + Aliro (n=7)** | **p-value** |
| --- | --- | --- | --- |
| **Body weight (g)** | 31.4 | 31.0 | 0.62 |
| **Total cholesterol (mg/dL)** | 627.2 ± 113.3 | 653.2 ± 121.8 | 0.68 |
| **LDL-cholesterol (mg/dL)** | 403.9 ± 99.2 | 401.0 ± 101.7 | 0.95 |
| **Triglycerides (mg/dL)** | 226.7 ± 64.5 | 297.4 ± 106.4 | 0.15 |
| **TNF-α (pg/mL)** | 4.4 ± 1.0 | 3.2 ± 0.5 | 0.04 |
| **IL-6 (pg/mL)** | 4.4 ± 2.4 | 2.4 ± 0.6 | 0.04 |

### 3.3 Alirocumab reduced plaque lesions and local inflammation in ApoE⁻/⁻ mice

We further investigated potential anti-inflammatory effects of alirocumab in the atherosclerotic plaque. As previously described^12,23^, ApoE⁻/⁻ mice exhibited prominent plaque lesions rich in lipid content at the aortic root (Figure 3A, top left). Importantly, alirocumab lessened plaque area (276 × 10^3^ µm^2^ vs. 655 × 10^3^ µm^2^; p<0.05) and its lipid accumulation (3.1 ± 0.5% vs. 5.1 ± 2.3%; p<0.01) (Figure 3A, top right). In addition, alirocumab diminished infiltration of CD68^+^-macrophages (2.3 ± 1.2% vs. 3.8 ± 0.8%; p<0.05) and CD80^+^-M1 macrophages (7.1 ± 2.8% vs.19.4 ± 7.0%; p<0.01) at the plaque lesion (Figure 3A, middle-bottom). Moreover, the TLR4 (2.3 ± 1.1% vs. 4.7 ± 1.1%; p<0.01), NLRP3 (2.2 ± 1.2% vs. 4.3 ± 1.2%; p<0.05) and caspase-1 (2.8 ± 1.1% vs. 5.3 ± 1.4%; p<0.01) content was also reduced after treatment (Figure 3B). Furthermore, in the descending aorta, the expression of major proinflammatory cytokines such as *CXCL10*, *TNF-α*, and *MCP-1* decreased after alirocumab, whereas anti-inflammatory cytokines (i.e., *TGF-β1*) and M2 macrophage membrane markers (i.e., *CD163* and *CD206*) were elevated (Figure 4).

**Figure 3.**
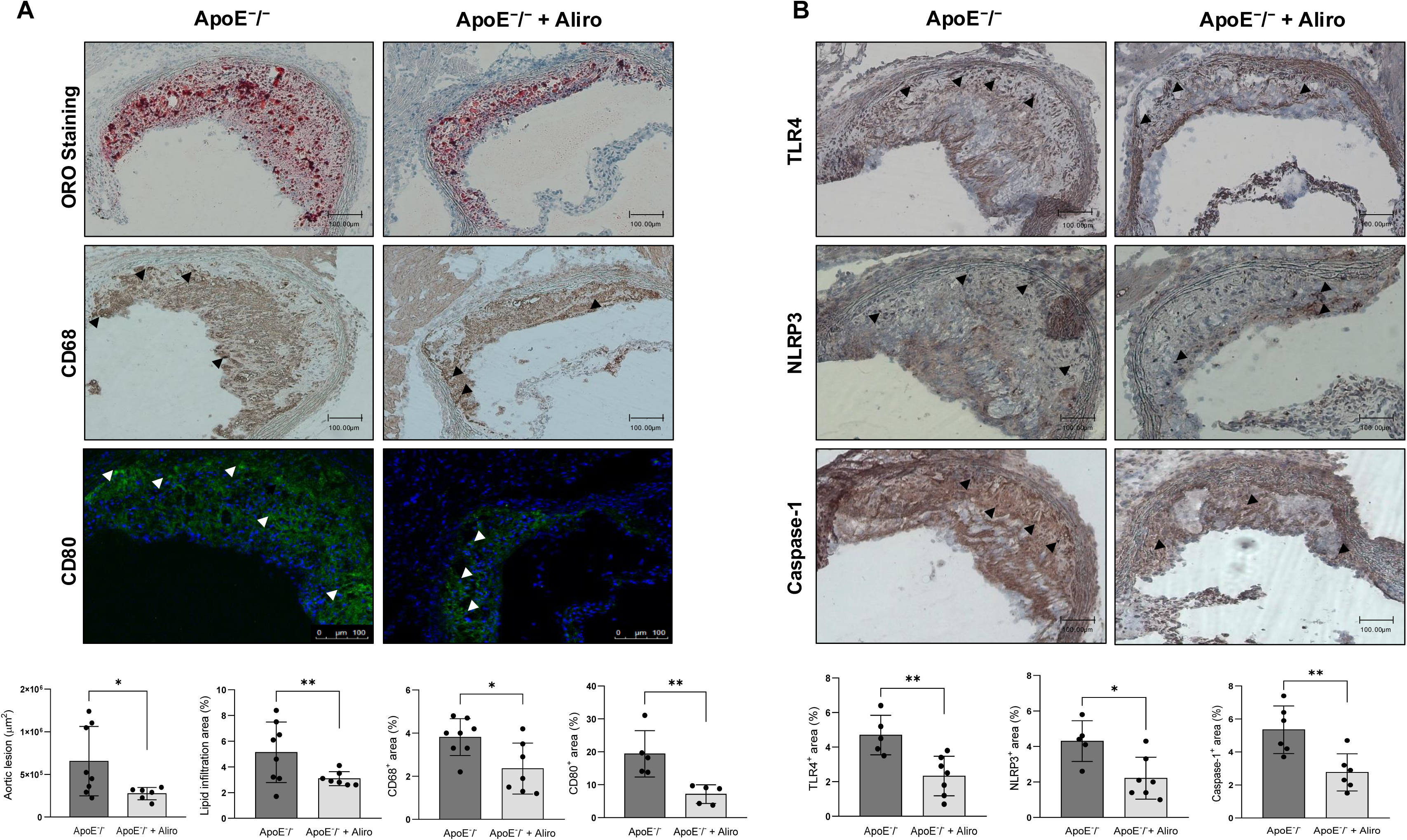
Alirocumab decreased lesion size, lipid accumulation and inflammation in atherosclerotic plaques from ApoE⁻/⁻ mice. **(A)** Representative images of the maximal atherosclerotic lesion at the aortic arch after staining with ORO or after incubation with anti-CD68 or -CD80 antibodies. Quantifications are expressed as lesion area and percentage of lipid accumulation (ORO-staining) or CD68^+^ and CD80^+^ macrophages in total intimal plaque area. Scale bar, 100 <u>µ</u>m. **(B)** Representative immunohistochemical staining of TLR4, NLRP3 and caspase-1 at maximal atherosclerotic lesions. Quantifications are expressed as percentage of total intimal plaque area. *p<0.05 and **p<0.01 vs. ApoE⁻/⁻ mice, by unpaired two-tailed Student’s *t*-test. N=5-9, per group. Scale bar, 100 <u>µ</u>m.

**Figure 4.**
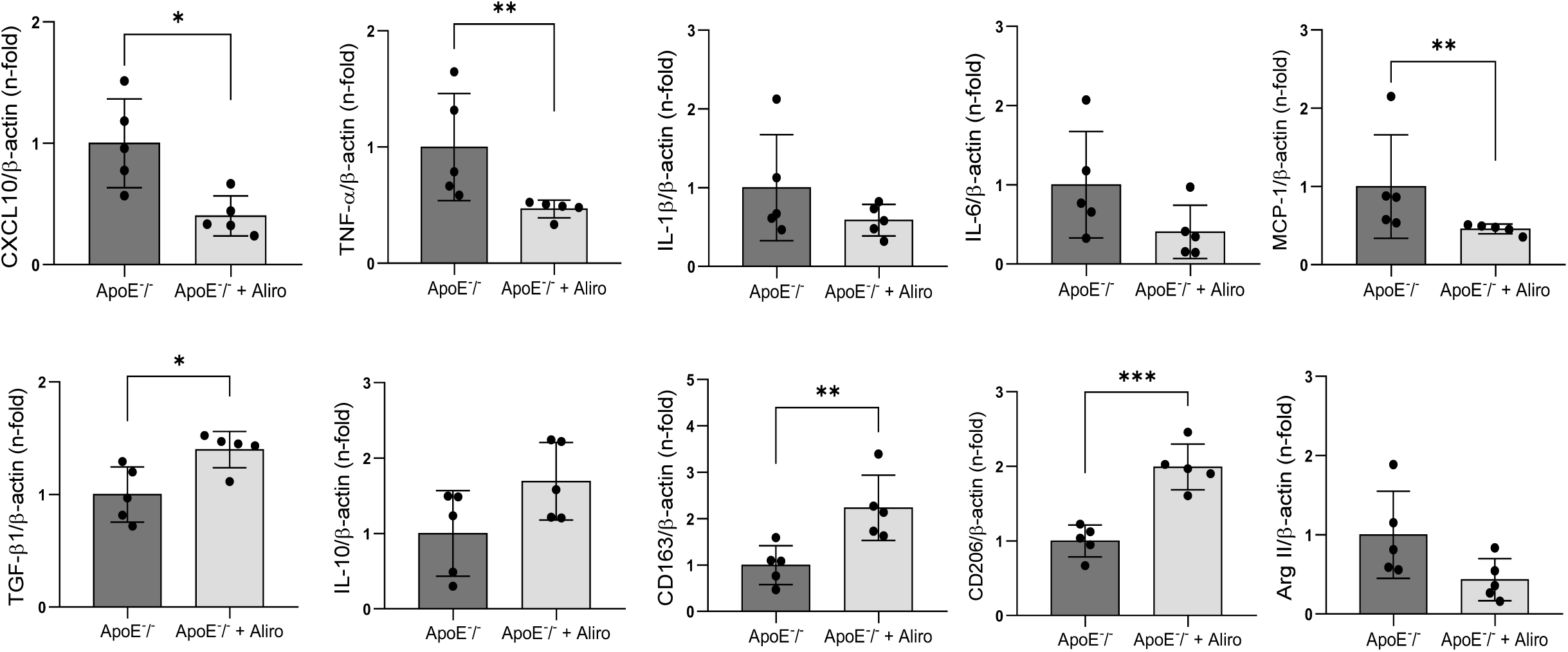
Alirocumab lessen the expression of proinflammatory cytokines while upregulating anti-inflammatory factors in ApoE⁻/⁻ mice. By qPCR, expression of proinflammatory *CXCL10, TNF-α, IL-1β, IL-6* and *MCP-1* cytokines, and anti-inflammatory *TGF-β1* and *IL-10* in descending aorta from ApoE⁻/⁻ mice with/without alirocumab treatment. Specific markers for M2 (*CD163, CD206)* and M1 (*Arg II*) were also evaluated. *p<0.05, **p<0.01, and ***p<0.001 vs. ApoE⁻/⁻ mice, by unpaired two-tailed Student’s *t*-test or U-Mann Whitney test depending on data distribution. N=5, per group.

### 3.4 PCSK9 upregulated inflammatory factors in cultured M1 macrophages

Next, we wondered whether PCSK9 might trigger direct proinflammatory actions on macrophages. Human THP-1 cells were polarized to M1 macrophages, as confirmed by elevation of the *CXCL9/CCL17* ratio (Suppl. Fig. 1C), and were incubated with recombinant PCSK9 for 24 hours. Remarkedly, the expression of proinflammatory cytokines including *TNF-α* (1.8 ± 0.5-fold vs. unstimulated M1 cells; p<0.05), *IL-1β* (2.8 ± 0.8-fold; p<0.05), *IL-6* (1.5 ± 0.1-fold; p<0.01), *CXCL9* (2.2 ± 0.5-fold; p<0.05), and *CXCL10* (3.7 ± 0.5-fold; p<0.05) was upregulated, in a similar manner than TNF-α (Figure 5, top). In contrast, PCSK9 downregulated anti-inflammatory cytokines such as *CCL17* (0.5 ± 0.1-fold vs. unstimulated M1 cells; p<0.01), *TGM2* (0.7 ± 0.2-fold; p<0.05), *IL-10* (0.8 ± 0.2-fold; p < 0.05), and *TGF-β1* (0.5 ± 0.2-fold; p<0.01) (Figure 5, bottom).

**Figure 5.**
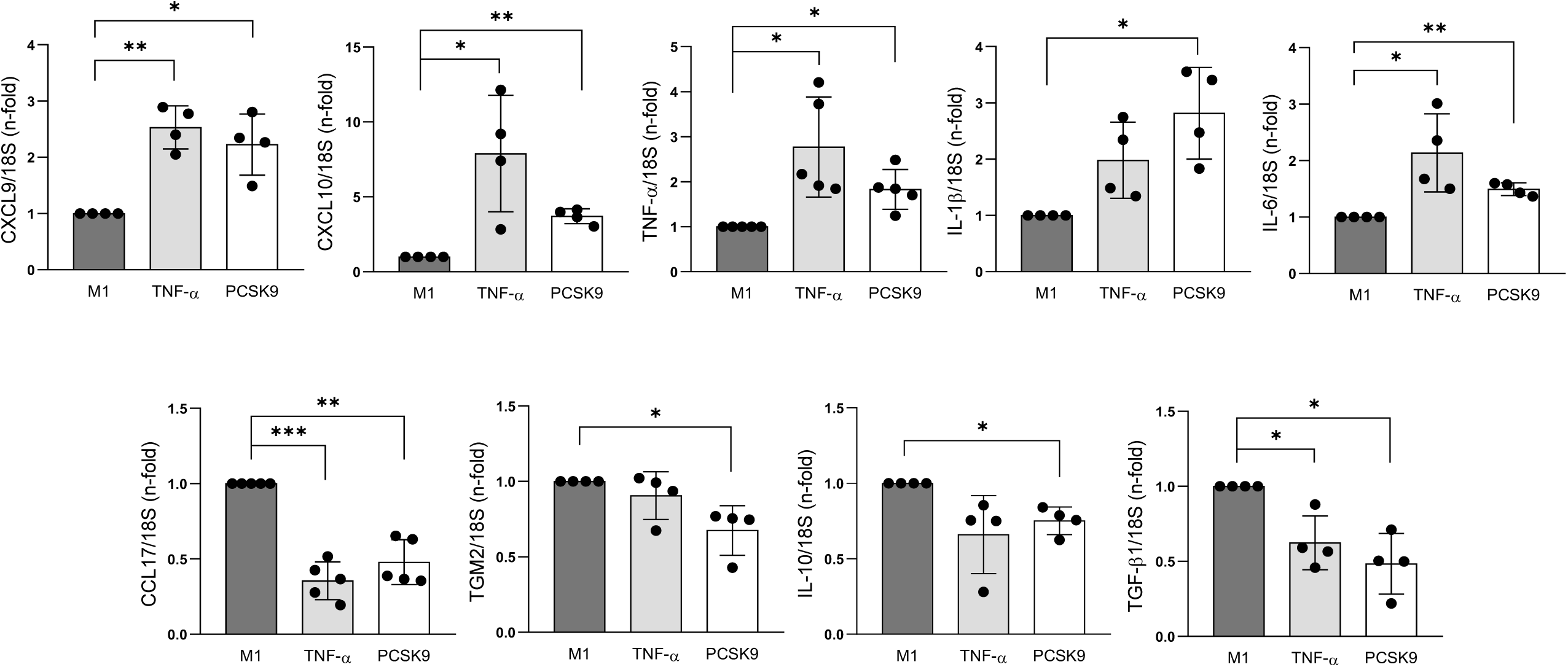
PCSK9 upregulated proinflammatory cytokines while decreasing anti-inflammatory factors in M1 macrophages. By qPCR, levels of proinflammatory *CXCL9*, *CXCL10, TNF-α, IL-1β* and *IL-6* and anti-inflammatory *CCL17, TGM2, IL-10* and *TGF-β1* cytokines in M1 macrophages stimulated with PCSK9. TNF-α was used as a positive control. *p<0.05, **p<0.01, and ***p<0.001 vs. unstimulated cells, by One-sample t test, control value set to 1. N=4-5 independent biological replicates.

In this sense, we also evaluated in M1-polarized macrophages the activation of major proinflammatory transcription factors. PCSK9 induced translocation of the NFκB p65 subunit to the nucleus after 3 hours of incubation, similarly to TNF-α (Suppl. Fig 2). In fact, pre-treatment with the pharmacological NFκB inhibitor, parthenolide, attenuated this effect (Figure 6A). Parthenolide also reduced IκBα phosphorylation, which anchorage p65 to the cytosol, and expression of a major component of the inflammasome complex, NLRP3 (Figure 6A). Indeed, preincubation with MCC-950, an inhibitor of inflammasome assembly, significantly lessened PCSK9-induced NLRP3 expression, but not p65 or IkBα (Figure 6B). Subsequently, MCC-950 also led to reduction in gasdermin-D cleavage, a downstream effector of caspase-1–mediated NLRP3 activation. Therefore, PCSK9 could promote direct proinflammatory actions in M1 macrophages through activation of NFκB and NLRP3-inflammasome.

**Figure 6:**
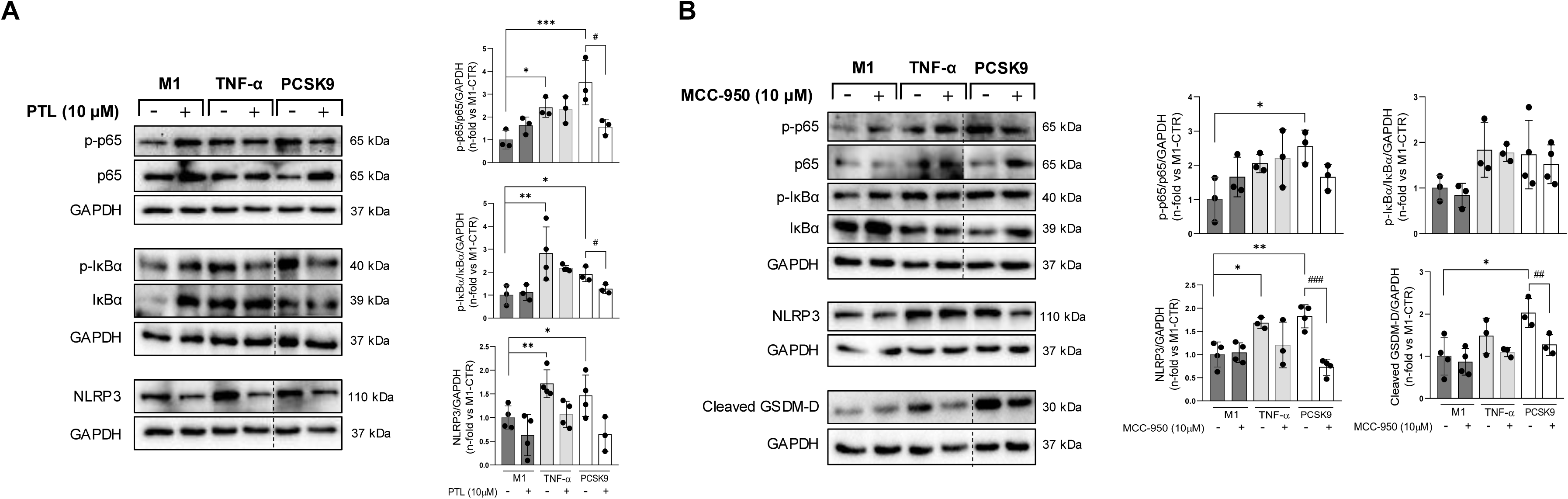
PCSK9 activated NFκB and NLRP3-inflammasome in M1 macrophages. **(A)** Phosphorylated and total p65 and IκBα were identified in PCSK9-stimulated M1 macrophages with/without pre-treatment of a NFκB inhibitor, parthenolide. Ratios of phosphorylated vs. non-phosphorylated were quantified. Bottom, NLRP3 was also detected and quantified. **(B)** NLRP3 and cleaved gasdermin-D were revealed after PCSK9 stimulation in M1 macrophages, with/without pre-treatment with a NLRP3 inhibitor (MCC-950). Phosphorylated and total p65 and IκBα were also uncovered, and the ratios of both isoforms were quantified. TNF-α was used as positive control. *p<0.05, **p<0.01, and ***p<0.001 vs. unstimulated cells, by one-way ANOVA followed by Dunnett’s post hoc test, and ^#^p>0.05, ^##^p>0.01, and ^###^p>0.001 vs. PCSK9, by one-way ANOVA with Tukey’s correction. We used at least three independent biological replicates.

### 3.5 TLR4, as a potential receptor for proinflammatory actions of PCSK9 in M1 macrophages

We then hypothesized that these direct PCSK9-proinflammatory actions might be mediated by pattern recognition receptors to translate danger signals from injured tissues to trigger inflammatory responses. Interestingly, in M1 macrophages stimulated with PCSK9, pre-treatment with a TLR4 antagonist, TAK-242 resulted in a marked lessening of p65 and IκBα phosphorylation, as well as a decline in NLRP3 and cleaved gasdermin-D (Figure 7). Importantly, these effects were confirmed by TLR4 gen silencing in PCSK9-stimulated M1 macrophages (Suppl. Fig. 3A). In turn, PCSK9 was able to increase TLR4 and its signalling adapter, MyD88 (1.9 ± 0.1 and 1.4 ± 0.1-fold; p<0.01, respectively, Suppl. Fig. 3B) suggesting that TLR4 may work as a critical downstream mediator of PCSK9 in M1 macrophages for proinflammatory outcomes.

**Figure 7.**
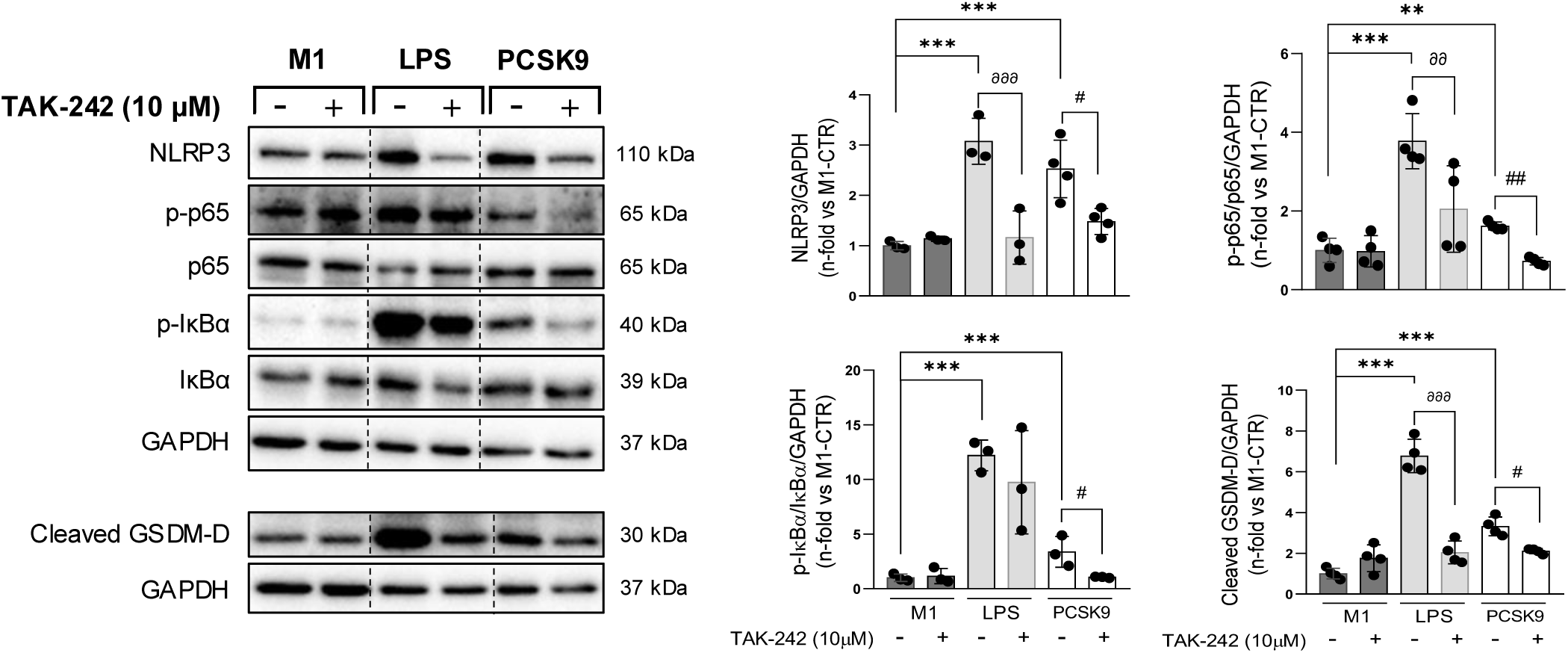
TLR4 inhibition dampened activation of the NFκB–NLRP3 axis upon PCSK9 stimulation. NLRP3 and cleaved gasdermin-D were identified after PCSK9 stimulation in M1 macrophages, with or without pre-treatment a TLR4 inhibitor, TAK-242. Phosphorylated and total p65 and IκBα were also recognized, and the ratios of both isoforms were calculated. LPS was used as positive control. *p<0.05, **p<0.01 and ***p<0.001 vs. unstimulated cells, by one-way ANOVA followed by Dunnett’s post hoc test. ^∂∂^p<0.01 and ^∂∂∂^p<0.001 vs. LPS, by one-way ANOVA with Tukey’s correction; and ^#^p>0.05 and ^##^p<0.01 vs. PCSK9, by one-way ANOVA with Tukey’s correction. We used at least three independent biological replicates.

### 3.6 Alirocumab attenuated PCSK9-induced proinflammatory responses in M1 macrophages

Finally, we evaluated whether alirocumab directly counteracts PCSK9-induced inflammation in M1 macrophages. As expected, alirocumab significantly reduced the expression of *CXCL9* (2.3 ± 0.2 vs. 1.5 ± 0.1-fold in unstimulated cells; p<0.001), *CXCL10* (1.5 ± 0.2 vs. 2.1 ± 0.3-fold; p<0.05), *TNF-α* (1.0 ± 0.1 vs. 2.2 ± 0.5-fold; p<0.001), *IL-1β* (0.9 ± 0.1 vs. 2.0 ± 0.4-fold; p<0.01), and *IL-6* (1.6 ± 0.3 vs. 3.0 ± 1.4-fold; p<0.05) promoted by PCSK9, while enhanced anti-inflammatory cytokines such as *CCL17* (0.9 ± 0.3 vs. 0.3 ± 0.1-fold; p<0.01), *IL-10* (0.9 ± 0.4 vs. 0.4 ± 0.1-fold; p<0.05), and *TGF-β1* (1.2 ± 0.8 vs. 0.8 ± 0.1-fold, p<0.01) (Suppl. Fig. 4A). Moreover, alirocumab significantly lessened phosphorylated IκBα and NLRP3 expression after PCSK9 exposure (Suppl. Fig. 4B). In fact, alirocumab was able to diminish TLR4 transcripts, and perhaps Myd88, after PCSK9 stimulation (Suppl. Fig. 3B). Altogether, PCSK9 could induce proinflammatory actions in M1 macrophages independently of LDL-C mitigation and via TLR4, NFκB and NLRP3 signalling.

## 4. Discussion

In large clinical trials like the ODYSSEY OUTCOMES, alirocumab demonstrated effective reduction of atherosclerotic plaques and cardiovascular events, but beyond lipid-lowering actions on LDL-C, alirocumab could have also exerted direct anti-inflammatory effects^25,26^. Herein, we demonstrated that alirocumab may attenuate progression of experimental atherosclerosis by controlling PCSK9-stimulated TLR4-NFκB-NLRP3 axis in M1 macrophages.

The notion that PCSK9 may influence macrophage plasticity was previously suggested in experimental infarcted hearts. The PCSK9 gene silencing caused a reduction in M1 macrophage content and an increase in M2 phenotype, which ameliorated inflammatory infiltration and infarct size^27^. Also, PCSK9-knockout mice displayed a decreased inflammatory response to LPS and PCSK9 pharmacological inhibition improved inflammation^28^. Interestingly, PCSK9 overexpression in ApoE⁻/⁻ mice led to infiltration of pro-inflammatory Ly6C^high^ monocytes into atherosclerotic plaques, independently of plasma lipid levels^29^. In this sense, we observed that PCSK9 plasma levels were associated with other proinflammatory factors such as hsCRP and FGF-23 in patients with ACS, even after adjustment by presence of previous coronary disease, dyslipemia, and statin use. Also, in mice, inhibition with alirocumab triggered a systemic anti-inflammatory phenotype characterized by a marked reduction in splenic M1 macrophages and elevation of reparative M2 cells, despite LDL-C levels were unaltered. Alirocumab also decreased plaque area, lipid accumulation and overall macrophage infiltration, preferentially limiting the M1 subset. These benefits were consistent with other reports in ApoE⁻/⁻ and LDLR⁻/⁻ models, where the size of aortic lesion, macrophage infiltration, and expression of inflammatory mediators (i.e., TNF-α, IL-1β, and MCP-1) were lessen without changes in plasma cholesterol^30,31,32^. Also, evolocumab triggered lipid-independent effects, including enhancement of autophagy and reduction in oxidative stress to improve plaque size and composition^33^. Therefore, inhibition of PCSK9 by neutralizing antibodies might lead to additional non-lipid-mediated protective effects that could complement the anti-atherogenic actions of these drugs or other treatments (i.e., statins)^9^.

Concerning to molecular mechanisms, it is being postulated that polarization of macrophages within the atherosclerotic plaque is a key determinant of its progression and stability. Pro-inflammatory M1 macrophages, characterized by high TLR4 expression, dominate the microenvironment of advanced plaques^34^. Much of the existing literature has focused on demonstrating that PCSK9 can initiate the polarization of naïve macrophages towards this M1 phenotype^35^. Thus, our study specifically investigated PCSK9 effects on M1-activated macrophages, reflecting the established inflammatory environment of atherosclerotic lesions^36^. In this line, we observed that PCSK9 could amplify the M1 phenotype, increasing the CXCL9/CCL17 ratio and secretion of pro-inflammatory cytokines (IL-1β, IL-6, TNF-α) via the TLR4-NFκB-NLRP3 pathway, similarly to TNF-α. This is consistent with previous studies in murine macrophages and endothelial cells where PCSK9 overexpression stimulated IκBα phosphorylation and p65 translocation^37^, while PCSK9 knockdown diminished TLR4 expression and NFκB activation in atherosclerotic plaque lesions. More recently, cyclase-associated protein-1 (CAP-1) has been identified as a high-affinity binding partner for PCSK9 in monocytes, acting upstream of TLR4^38^. Thus, TLR4 can be an indispensable mediator to transduce PCSK9 signaling toward NFκB and downstream inflammasome activation. Activation of NFκB was previously suggested as an associated factor of PCSK9-stimulated monocytes^39,40^, but the inflammasome could strengthen the proinflammatory signalling. Particularly, the NLRP3 inflammasome can be a critical innate immune sensor that influence atherosclerosis by detecting endogenous danger signals^41,42^. Its activation induces caspase-1–mediated IL-1β maturation and gasdermin-D cleavage, triggering pyroptosis and intensifying plaque inflammation^43^. Also, patients treated with PCSK9 inhibitors displayed lower NLRP3 and active caspase-1 expression within carotid plaques compared to patients receiving other lipid-lowering therapies, even after adjustment by LDL-C levels^44^.

Although these findings may have considerable interest, our study has some limitations. Firstly, translating these findings to humans remains challenging since alirocumab is primarily a lipid-lowering therapy and thus, its immunomodulatory effects should be considered instead additional or complementary. Also, although our data clearly show increased gasdermin-D/Caspase-1 upon PCSK9 stimulation, additional studies are warranted to delineate the extent and regulation of pyroptotic signalling. Finally, the effect of PCSK9 on other immune cells such as neutrophil or monocytes-derived subtypes (i.e., M0 and M2 macrophages) may add interesting information of PCSK9 on the inflammatory response after atherosclerosis. In short, PCSK9 could activate the TLR4-NFκB-NLRP3 axe to produce lipid-independent inflammatory actions on vasculature and may behave as a dual factor reducing lipid clearance. Thus, targeting this pathway with PCSK9 inhibitors like alirocumab might provide therapeutic benefits on atherosclerosis and other inflammatory-based cardiovascular diseases.

## 5. Clinical implications

These findings highlight a novel lipid-independent mechanism by which PCSK9 contributes to vascular inflammation through activation of the TLR4–NFκB–NLRP3 axe in M1 macrophages. Thus, beyond lowering LDL-C, PCSK9 inhibitors such as alirocumab could mitigate macrophage-driven inflammation and reduce atherosclerotic plaque progression. This dual role may position alirocumab as a promising therapeutic approach to stabilize atherosclerotic plaques and improve cardiovascular diseases such as acute coronary syndrome.

## 6. Acknowledgments

The authors would like to thank Ghassik Munther for her contribution to this work as part of her undergraduate thesis.

## 9. Authors contribution

CE participated in the conception and design of the study, performed all in vitro and preclinical experiments. MS-C, MR-C, MK, and IH-DR contributed to the development of the in vivo experiments. AO-H and DG-G participated in the flow cytometry analysis. JL-C contributed to clinical study. CG-G provided the in vivo model and contributed to the experimental design. JE provided supervision and critical manuscript revision. JT and OL are the principal investigators of the study and were responsible for the overall supervision.

## 10. Data availability

All data supporting this study are available within the Supplementary Material. Additional data related to the observational study are available upon reasonable request.

## Non-standard Abbreviations and Acronyms

CCL17: C-C motif chemokine ligand 17
CXCL9: C-X-C motif chemokine ligand 9
CXCL10: C-X-C motif chemokine ligand 10
FGF-23: Fibroblast growth factor-23
hsCRP: High-sensitive C-reactive protein
IL-1β: C-reactive protein
IL-6: Interleukin-6
IL-10: Interleukin-10
LDL-C: Low-density lipoprotein cholesterol
MCP-1: Monocyte chemoattractant protein-1
M1: Classically activated macrophage (pro-inflammatory phenotype)
M2: Alternatively activated macrophage (anti-inflammatory phenotype)
NFκB: Nuclear factor kappa-light-chain-enhancer of activated B cells
NLRP3: NOD-like receptor family pyrin domain-containing 3
PCSK9: Pro-protein convertase subtilisin/kexin type 9
TGF-β1: Transforming growth factor beta 1
TGM2: Transglutaminase 2
TLR: Toll-like receptor
TNF-α: Tumor necrosis factor Alpha

---

**Supplementary Figure 1. (A) Gene silencing of TLR4 in M1 macrophages.** Transcript and protein levels of PCSK9 after pre-treatment with siRNA-TLR4. *p<0.05 and ***p<0.001 vs. scramble (by one-sample t test (mRNA) or by unpaired two-tailed Student’s *t*-test (protein)). N=3, independent biological replicates. **(B)** Plasma and liver PCSK9 levels in the experimental model ***p<0.001 vs. ApoE⁻/⁻ mice (by unpaired two-tailed Student’s *t*-test). N=7-9, per group. **(C) Polarization of M0 into M1 macrophages.** Relative expression of the ratio of *CXCL9* and *CCL17* after 48h incubation with polarization medium. *p<0.05 vs. M0 by One-sample t test, control value set to 1. N=4, independent biological replicates.

**Supplementary Figure 2. PCSK9 promoted nuclear translocation of the NFκB-p65 subunit in M1 macrophages.** Representative immunofluorescence images of M1 macrophages showing NFκB-p65 (green) and nuclei (blue) staining after PCSK9 (or TNF-α) stimuli. Right, quantification of positive co-localization of TNF-α and PCSK9 with nuclear staining (DAPI). ***p<0.001 vs. unstimulated cells, by one-way ANOVA followed by Dunnett’s post hoc test. N=3, independent biological replicates. Scale bar, 75 µm.

**Supplementary Figure 3. (A) Participation of TLR4 in the PCSK9-induced NFκB and NLRP3 activation.** NLRP3 and cleaved gasdermin-D were identified after PCSK9 stimulation in M1 macrophages, with/without pre-treatment of siRNA-TLR4. Phosphorylated and total p65 and IκBα were also identified, and the ratios of both isoforms were calculated. LPS was used as positive control. **(B) Implication of a TLR4 mediator in PCSK9 inhibition.** Myd88 expression was analyzed after PCSK9 with/without pre-treatment with alirocumab. *p<0.05, **p<0.01, and ***p<0.001 vs. unstimulated cells by one-way ANOVA followed by Dunnett’s post hoc test for protein expression vs. unstimulated cells; ^∂^p<0.05 and ^∂∂^p<0.01 vs. LPS and ^#^p>0.05 vs. PCSK9, by one-way ANOVA with Tukey’s correction. (N= 3-4)

**Supplementary Figure 4. Alirocumab attenuated pro-inflammatory genes and mediators induced by PCSK9 in M1 macrophages**. **(A)** By qPCR, levels of proinflammatory (C*XCL9, CXCL10, TNF-α, IL-1β* and *IL-6* and anti-inflammatory (*CCL17, TGM2, IL-10,* and *TGF-β1*) cytokines in M1 macrophages stimulated with PCSK9 with/without alirocumab. *p<0.05, **p<0.01, ***p<0.001 vs. unstimulated cells, by one-sample t-test; ^#^p>0.05, ^##^p>0.01, ^###^p<0.001 vs. PCSK9, by by t-test. N=4 independent biological replicates. **(B)** Phosphorylated and total p65 and IκBα were identified in PCSK9-stimulated M1 macrophages with/without alirocumab. Ratios of phosphorylated vs. non-phosphorylated isoforms were quantified. Bottom, NLRP3 was also detected and quantified. *p<0.05 vs. unstimulated cells by one-way ANOVA followed by Dunnett’s post hoc test; ^#^p>0.05 and ^##^ p> 0.01 vs. PCSK9 by one-way ANOVA with Tukey’s correction. We used at least three independent biological replicates.

